# First complete genome characterization of an Indian pigeon pox virus directly from a clinical sample

**DOI:** 10.1101/2023.11.19.567771

**Authors:** Basanta Pravas Sahu, Subhasmita Panda, Ravi Raj Singh, Subrat Kumar Swain, Niranjana Sahoo, Anjan Kumar Sahoo, Debasis Nayak

## Abstract

Avian pox disease is a highly contagious infection caused by pox virus and has serious consequences on avian species with regards to economic and conservation aspects. This viral genus named as Avipox virus (APV) that infects nearly 300 bird species and lack of enough complete genome information creates hindrance to infer this virus biology. Thus in this study, we have revealed the first complete genome of an Indian pigeon pox virus that belongs to the genus APV followed by comparative genomics analysis. The entire genome of present isolate (PPV/Pur-Od-4b/01/Ind) having 280058 bp nucleotide sequences with the GC content 29.51%. The unique feature of this complete genome revealed the presence of 270 open reading frames (ORFs) circumscribed by inverted terminal repeats (ITRs) of 4,689 bp at each end and lack of recombination events. The concatenated amino acid phylogenetic tree deciphered the present isolate closely related with Feral Pigeon pox virus derived from Africa. The molecular markers, such as microsatellites were ubiquitously distributed throughout the genome and more prevalent within the functional genes.

## 1 Introduction

Prevalence of pox virus emerged as a great threat to both domestic as well as free ranging wild birds irrespective to their age and sex due to its exorbitant contagious property, which influence both the economical and conservation aspects (Murphy et al., 1999). Pox like infections have been reported among wild and domestic bird species globally derived from 20 orders which includes 76 families having 329 avian species across the globe, caused by a group of viruses named as family avipox viruses (APVs) (Carulei et al., 2017). Although antigenetically and immunologically these viruses were distinct from one another, the cross relation within the broad host range arose the complications for viral strain demarcation. However, the virus nomenclature usually determined according to the host, from which the virus derived. Few of the examples included fowlpox virus (FWPV), turkeypox virus (TKPV), pigeonpox virus (PGPV), etc. APVs, present throughout the globe that cause highly transmissible avian pox disease which spread by viral vectors such as arthropods like mosquitoes, mites, flies, from diseased birds to healthy one through contaminated air, food or water (Adebajo et al., 2012). The severity of the infection generally depends upon two forms the disease. The first one is mild cutaneous lesions (dry form), rarely fatal in comparison to the second form such as diphtheric (wet). The secondary bacterial and fungal infection increases the rate of morbidity and mortality (up to 60%) by forming the painful lessons around the eye, beak respiratory track that interfere with eating, respiration and vision (Skinner, 2008). APVs belongs to DNA viruses having enveloped lipid membrane, oval-shaped with the particle size 270 × 350 nm and centrally located conserves regions and variable terminal inverted repeats (Afonso et al., 2000; Weli & Tryland, 2011). Due to its too large genome size with stable central region and host range-restricted to avian species increase the efficacy of some of the virus like (FPV) utilized to construct recombinant vaccines for foot and mouth disease virus (Zheng et al., 2006)), Influenza (Taylor et al., 1988) and HIV (Jiang et al., 2005).

Although a number of reports suggest the emergence of avipoxvirus worldwide including India, causing infection in fowl, turkey, pigeon, duck and quail with high morbidity and mortality, still a long way to move to decipher its genome complexity (Abdallah & Hassanin 2013; Gyuranecz et al., 2013; Sharma et al., 2019; Sahu et al., 2020). The lack of enough surveillance as well as vaccine failure in India indicates the re-emergence of the viral species. Thus, we have taken the pigeonpox virus into our consideration as it is an infectious agent to Indian pigeons, which has been utilized as a food source, sport (racing), show animals and household pets. For the first time we deciphered the complete genome sequence of a pigeon pox virus (PPV) of Indian origin through next generation sequencing platform (NGS) directly from clinical specimen and utilize comparative genomics approach to reveal more about this virus biology.

## 2 Materials and methods

To decipher the complete genome sequence of Indian PPV, we have utilized the isolates PPV/Pur-Od-4b/01/Ind previously identified from our lab (Sahu, Majee et al. 2020). The extraction of nucleic acid from infected scab samples was performed by using DNeasy blood and tissue purification Kits (QIAGEN, USA) with some modification established by Sarker et al., 2017 (Sarker, Das et al. 2017). The quality and quantity of DNA checked with 1% agarose gel followed by nanodrop estimation. Finally, 250 ng of genomic DNA send for library construction followed by paired-end sequencing chemistry on NextSeq500 sequencing platform.

The raw data generated through NGS platform further cleaned by Trimmomatic v0.38 to eliminate noisy or ambiguous reads, adapter and host sequence (Bolger et al., 2014). The quality reads further mapped to reference sequence utilizing BWA MEM software (version 0.7.17) (Li & Durbin, 2009). The extraction of consensus reads Along with Denovo assembly was performed using SAM tools (Li and Durbin, 2009). GATU was used to identify possible Open reading frame (ORFs) as well as genome annotation (Tcherepanov et al., 2006). The presences of possible genes or ORFs were identified depending upon the significance sequence similarity with known cellular or viral proteins. To infer evolutionary relationship of present virus with other APVs, a phylogenetic tree was constructed using concatenated amino acid sequences of the nine poxvirus core proteins such as RNA polymerase subunit RPO132, RNA polymerase subunit RPO147,mRNA cappin<u>g</u> enzyme large subunit, RNA polymerase-associated protein RAP94, virion core protein P4a, virion core protein P4b, early transcription factor large subunit VETFL, NTPase and DNA polymerase by following the method recently used to decipher the evolutionary relationship among APVs (Sarker et al., 2017). In this method, the selected sequences were aligned using the MAFTT followed by GTR substitution model with 1000 bootstrap replicates in MEGAX (Kumar et al., 2018). Furthermore recombination events were checked by using the RDP, GENECONV, Bootscan, MaxChi, Chimaera, Siscan, PhylPro, LARD and 3Seq methods of RDP4 program (Martin et al., 2015) within APVs with respect to present isolate. Parameters having a significant p-value within three of the methods referred as a possible recombination event. A genome wide scanning of perfect mono to hexa as well as compound microsatellites conducted using Krait software (Du et al., 2018). For the identification of perfect microsatellite the parameters such as: type of repeat: perfect; repeat size: all; minimum repeat number: 6, 3, 3, 3, 3, 3 for mono-hexa repeats, respectively were set. Maximum distance (dMax) allowed between any two microsatellites was 10 nucleotide. The rest of the parameters were kept as default to reveal the composition of simple SSRs and compound SSR (cSSRs).

## 3 Results and discussion

The NextSeq 500 NGS generated a total of **∼**2.72 GB of quality data with 9,541,974 numbers of qualities reads. The previously published complete genome isolate pigeonpox virus FeP2 (Acc. no-NC_024447) used as the reference genome, to which the sequence generated through NGS mapped and assembled. The assembled genome of this novel virus exhibited 280058 bp in length. Then the assembled genome annotated through sequin and submitted to GenBank to get the accession number ON375849. A lower percentage of GC content (29.5) was observed in this newly assembled genome, in comparison to fowlpox virus (30.83), shearwater pox virus (30.23), canarypox virus (30.37), turkeypox virus (29.78) but similar to other APV such aspigeonpox virus, flamingo pox virus, magpiepox virus and penguinpox virus. The extreme left nucleotide was considered as base 1, and the beginning region of the Inverted Terminal Repeats (ITRs), which consists of total length 4689 bp. The ITR spanned throughout PPV-001 to PPV-004a and PPV_256 toPPV_259. Using NCBI’s The ORF Finder tool of NCBI revealed the presence of 270 ORFs, from which 10 have been annotated as truncated and 34 fragmented genes. Among those genes, 112 genes orientated positively, and rest 158 have a negative orientation (Figure 1). The comparison of both predicted nucleotide (BLASTN) as well as proteins (BLASTP) with the other reference pigeonpox resulted 99% to 100%similarity. In the present isolate, we observed maximum mutation within hypothetical protein followed by ankyrin repeat protein. These mutations might be the major concern for the heterogeneity and virulence of the respective pathogen. The concatenated phylogenetic tree suggested the present PPV isolate shared a close relationship with FeP2 strain (Acc. No-NC_024447) (Fig. 1). Interesting, we did not observe any recombination events with respect to our isolate. However this kind of ambiguity was noticed within pox virus during previous study (Sarker et al., 2021).

**Figure 1:**
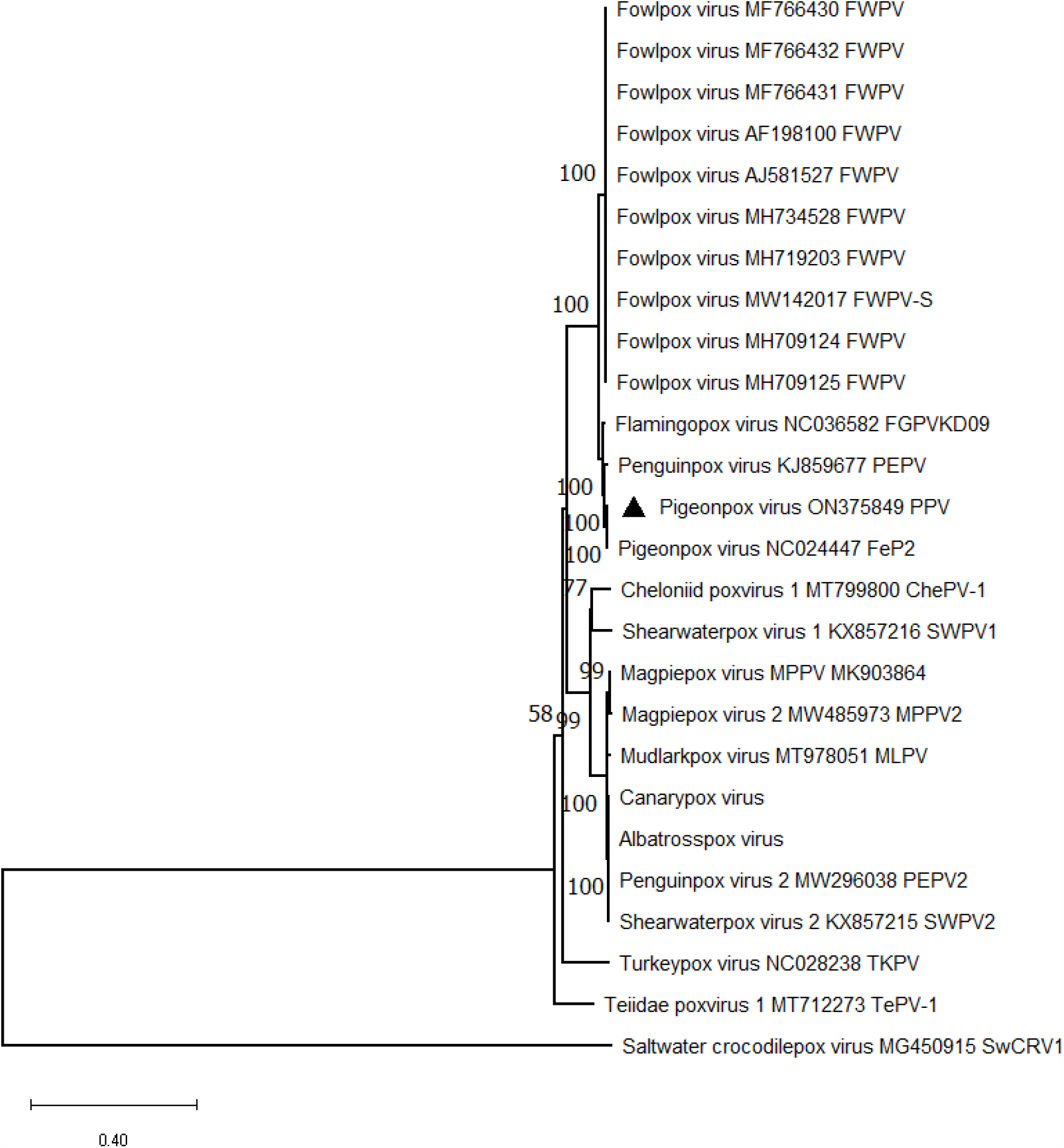
A concatenated phylogenetic tree deciphering the relationship between PPV/Pur-Od-4b/01/Ind and other chordopox viruses. To reveal the evolutionary relationship of present virus with other APVs, a phylogenetic tree was constructed using concatenated amino acid sequences of the nine poxvirus core proteins such as RNA polymerase subunit RPO132, RNA polymerase subunit RPO147, mRNA capping enzyme large subunit, RNA polymerase-associated protein RAP94, virion core protein P4a, virion core protein P4b, early transcription factor large subunit VETFL, NTPase and DNA polymerase followed by MAFFT alignment and GTR substitution model with 1000 bootstrap replicates in MEGAX.

Our study revealed 1988 and 153 numbers of SSRs and cSSR, scattered throughout the PPV genome. Dinucleotide repeats were the most abundant (55.18%) in the genome, followed by a mononucleotide (32.8%), trinucleotide repeats (11.12%), tetranucleotide (0.65%) and pentanucleotide (0.2%). Hexanucleotide repeats were observed at least in number and represented at least 0.05% within the PPV/Pur-Od-4b/01/Ind genome, respectively (Fig. 2). The SSRs were found in functional protein and hypothetical protein occupied 70% and 5%, respectively. In the case of cSSR, 69% and 31% of microsatellite motifs were distributed within coding or non-coding regions. Functional and hypothetical proteins occupied 67% and 21%, respectively. The percentage of individual microsatellites being part of compound microsatellite (cSSR%) was 16.3%. The overall frequency of mononucleotide repeats A/A (52.14%) most prevalent than poly G/C (0.76%), dinucleotide repeat motif AT/TA (43.84%) were most abundant over AT/AT (38.83%), AG/TC (3%), AG/AG (2.82%), AC/GT (2.5%) and AC/CA (2%), AC/TG (1.73) respectively. Microsatellites have a higher polymorphism, which was utilized for strain identification and evolutionary analysis in several viruses named as herpes simplex virus (Burrel et al., 2013) and adenovirus (Houng et al., 2009). However a recent study revealed the correlation between microsatellites with recombination hot spot (George et al., 2015). Thus, it would be interesting to reveal the role of abundant number of microsatellites present within the PPV/Pur-Od-4b/01/Ind genome in future research.

**Figure 2:**
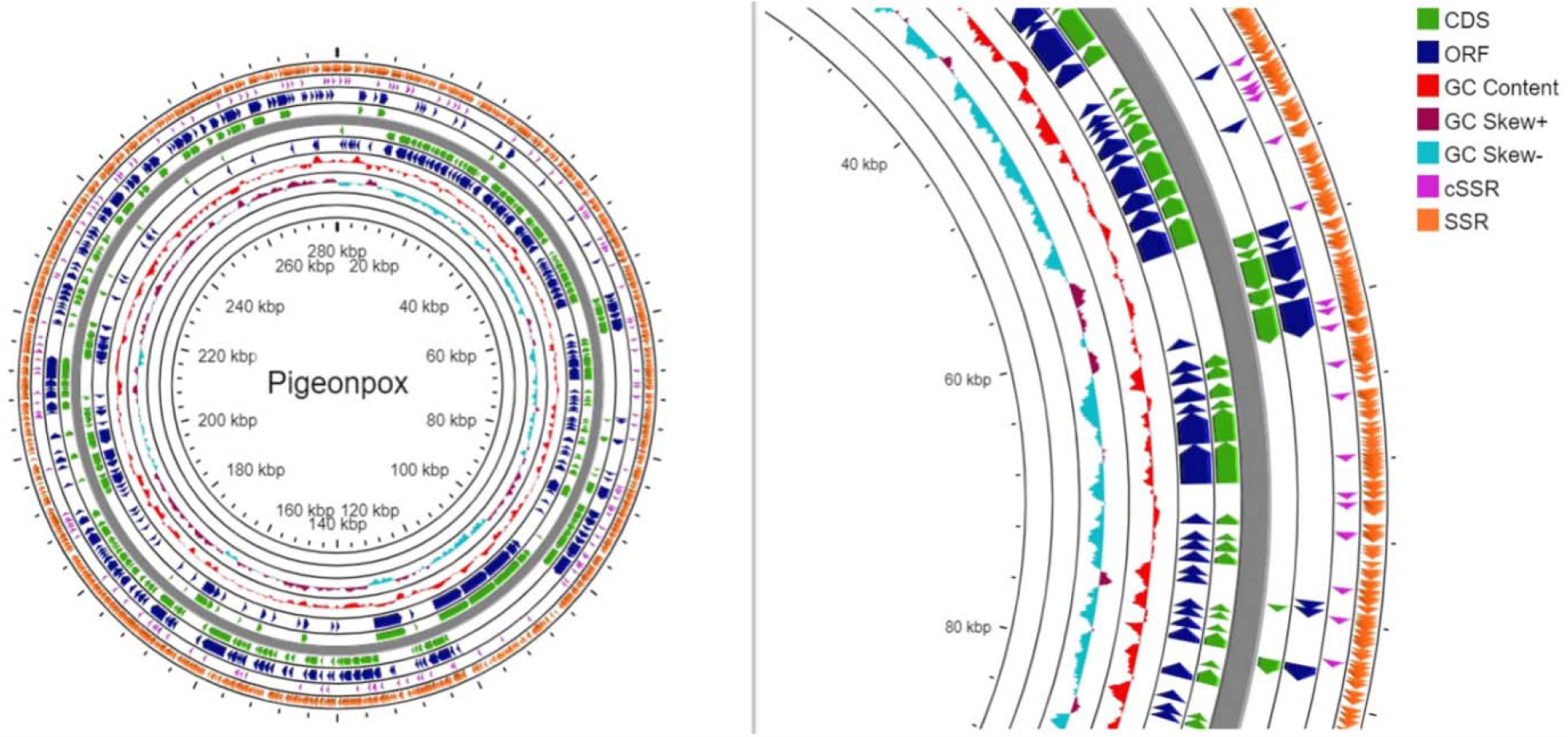
Circos map depicting complete genome of the PPV/Pur-Od-4b/01/Ind isolate. **Figure** representing all the coding sequences followed by repeat regions and open reading frame in different colors from inner to outer track: GC skew (maroon and sky blue), GC content (red), ORF (blue), CDS (green), cSSR (pink) and the outermost region represent the SSR (orange).

So far, there are no clear patterns regarding species-specificity in the APVs. Many bird species experience life-long immunity if the immune system is not weakened or the birds are not infected by different strains (Winterfield et al., 1985). Stressful conditions, poor nutrition, environmental contamination and ill health may contribute to the severity of such lesions. Very little is known about the incidence, morbidity or mortality rates of poxvirus infections in wild birds. Thus the present study will enrich the genomic resource as there is only one complete genome present in GenBank. Moreover this data set may help to understand the virus followed by development of efficient prevention strategy to limit the virulence.

## DATA AVAILIBILITY STATEMENNT

The complete genome sequence of present study is openly available in the GenBank reposatory (https://www.ncbi.nlm.nih.gov/genbank/) with the accession number ON375849.

## AUTHOR CONTRIBUTIONS

BPS and DN contributed to the conceptualization of this study. SP, SKS, carried out the genomic DNA isolation and figure construction. NS, AKS collected the samples. BPS and DN compiled the results and prepared the first draft. SP, SKS, NS and AKS supervised the study and edited the final draft. All authors read and approved the final manuscript.

## ACKNOWLEDGEMENTS

BPS is thankful to the University Grant Commission (UGC, Govt. India) for providing Ph.D. fellowship as financial support.

## Ethics statement

Filed veterinarians collected clinical samples as a routine practice from symptomatic birds. The guideline (clause 3.2.2) set by the Committee for Control and Supervision of Experiments on Animals (CPCSEA), Ministry of Environment and forest, Govt. of India was followed during the sample collection procedure.

## Conflict of Interest Statement

The authors declare no conflict of interest.

## Funding

BPS is supported by University Grant Commission (UGC, India) fellowship for his doctoral studies.

## Notes

### Competing Interest Statement

The authors have declared no competing interest.

